# Pangenomic genotyping with the marker array

**DOI:** 10.1101/2022.05.19.492566

**Authors:** Taher Mun, Naga Sai Kavya Vaddadi, Ben Langmead

**Affiliations:** Johns Hopkins University, USA, Illumina, USA; Johns Hopkins University, USA

**Author notes:** **Funding** *Taher Mun*: R01HG011392 and R35GM139602 to BL. Also, NSF-135491. *Naga Sai Kavya Vaddadi*: R01HG011392 and R35GM139602 to BL *Ben Langmead*: R01HG011392 and R35GM139602 to BL. Also, NSF-135491.

**Keywords:** Sequence alignment, indexing genotyping

## Abstract

We present a new method and software tool called `rowbowt` that applies a pangenome index to the problem of inferring genotypes from short-read sequencing data. The method uses a novel indexing structure called the marker array. Using the marker array, we can genotype variants with respect from large panels like the 1000 Genomes Project while avoiding the reference bias that results when aligning to a single linear reference. `rowbowt` can infer accurate genotypes in less time and memory compared to existing graph-based methods. The method is implemented in the open source software tool `rowbowt` available at https://github.com/alshai/rowbowt.

## 1 Introduction

Given DNA sequencing reads from a donor individual, genotyping is the task of determining which alleles the individual has at polymorphic sites. Genotyping from sequencing data, sometimes using low-coverage sequencing data together with genotype imputation, is a common task in human genetics [10] and agriculture [18]. In contrast to variant calling, genotyping is performed with respect to a catalog of known polymorphic sites. For instance, genotyping of a human can be performed with respect to the 1000 Genomes Project call set, which catalogs positions, alleles and allele frequencies for tens of millions of sites [2].

Many existing genotypers start by aligning reads to a single linear reference genome, e.g. the human GRCh38 reference [26]. Because this reference is simply one example of an individual’s genome, genotyping is subject to reference bias, the tendency to make mistakes in places where the donor differs genetically from the reference. This was shown in studies of archaic hominids [17], HLA genotypes [4] and structural variants [28]. A similar bias exists for methods that extract polymorphic sites along with genomic context, and search for these sequences in the reads [11, 27]. In particular, the bias remains if the flanking sequences are extracted from the reference and so contain only reference alleles.

Reference bias can be avoided by using a *pangenome* reference instead of a single linear reference. A pangenome can take various forms; it can be (a) a generating graph for combinations of alleles, (b) a small collection of linear references indexed separately, or (c) a larger collection of linear reference indexed together in a compressed way. Pangenome graphs (option a) and small collections of linear references (option b) have been studied in recent literature [23, 8, 15, 7]. Variant graphs are effective for genotyping, but have drawbacks in this context. First, haplotype information might be removed when adding variants to the graph, or might be included in the graph but not considered during the read mapping process. This can cause tools like Bayestyper [29] to consider many extraneous haplotype paths through the graph during genotyping, increasing running time. Second, variant graphs can grow exponentially – in terms of the number of paths through the graph – as variants are added, leading to a rapid increase in resource usage and likelihood of ambiguous alignments.

We sought a way to avoid reference bias by indexing and querying many linear references at once while keeping index size and query time low. Such an approach can take full advantage of linkage disequilibrium information in the panel, allowing no recombination events except those occurring in the panel. This avoids mapping ambiguity from spurious recombination events between polymorphic sites [23].

We propose a new structure called the *marker array* that replaces the suffix-array-sample component of the *r*-index with a structure tailored to the problem of collecting genotype evidence. Here we describe the marker array structure in detail. We compare its space usage and query time to those of the standard *r*-index and explore how accurately both structures are able to capture markers from a sequencing dataset. Finally, we benchmark it using real whole-genome human sequencing data and compare it to existing genotyping tools in terms of both genotyping accuracy and computational efficiency.

## 2 Background

### *r*-index

The *r*-index [14] is a compressed repeat-aware text index that scales with the non-redundant content of a sequence collection. It uses *O*(*r*) space where *r* is the number of same-character *runs* in the Burrows-Wheeler Transform (BWT) of the input text. Past work shows that the *r*-index can efficiently index collections of long-read-derived human genome assemblies [19] as well as large collections of bacterial genomes [1].

While the main contribution of the *r*-index was its strategy for storing and using a sample of the suffix array [14], even this sample is large in practice. We propose a new *marker array* structure that replaces the suffix array while retaining its ability to deduce when a read-to-pangenome match provides evidence for a particular allele at a polymorphic site. The design of the marker array flows from three observations. First, we can save space by storing auxiliary information about polymorphic sites (“markers”) only at the sites themselves. There are often far fewer sites harboring polymorphism than there are BWT runs. Second, pangenome suffixes starting with the same allele tend to group together in the suffix array, which can be exploited to compress the marker array structure. Third, while a suffix array entry is an offset into the pangenome requiring *O*(log *n*) bits, a marker need only distinguish markers and alleles, and so requires just *O*(log *M*) bits where M is the number of polymorphic sites.

## 3 Methods

### Preliminaries

Consider a string *S* of length *n* from ordered alphabet Σ, with operator ≺ denoting lexico-graphical order. Assume *S*’s last character is lexicographically less than the others. Let *F* be an array of *S*’s characters sorted lexicographically by the suffixes starting at those characters, and let *L* be an array of *S*’s characters sorted lexicographically by the suffixes starting immediately after them. The list *L* is the Burrows-Wheeler Transform [5] of *S*, abbreviated BWT.

The `BWT` can function as an *index* of *S* [13]. Given a pattern *P* of length *m* < *n*, we seek the number and location of all occurrences of *P* in *S*. If we know the range `BWT`(*S*)[*i*..*j*] occupied by characters immediately preceding occurrences of a pattern *Q* in *S*, we can compute the range `BWT`(*S*)[*i*′..*j*′] containing characters immediately preceding occurrences of *cQ* in *S*, for any character *c* ∈ Σ, since

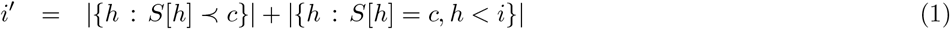

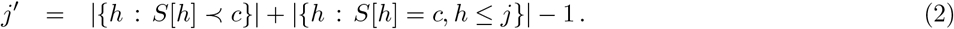

The FM Index is a collection of data structures for executing such queries efficiently. It consists of an array *C* storing |{*h* : *S*[*h*] ≺ *c*}| for each character *c*, plus a rank data structure for `BWT`(*S*), e.g. a wavelet tree, that can quickly tally the occurrences of a character *c* up to a position of `BWT`. To locate the offsets of occurrences of *P* in *S*, the FM-index can additionally include some form of *S*’s suffix array. The suffix array `SA` is an array parallel to *F* containing the offsets of the characters in *F*. To save space, the FM-index typically keeps only a sample of `SA`, e.g. a subset spaced regularly across `SA` or across *S*.

Let *T* = {*T*_0_, *T*_1_,…, *T_n_*} be a collection of *n* similar texts where *T*_0_ is the *reference sequence*, and *T*_1_, …, *T_n_* are *alternative sequences*. In the scenarios studied here, a *T_i_* represents a human haplotype sequence, with all chromosomes concatenated, and *T*_0_ represents the GRCh38 primary assembly of the human genome. Each *T_i_* with *i* > 0 is an alternate haplotype taken either from the 1000 Genomes project call set [2] or from the HGSVC project [6, 12], each with chromosomes concatenated in the same order as *T*_0_’s. We use the terms “haplotype” and “genome” interchangeably here.

We assume that all the *T_i_*’s are interrelated through a multiple alignment, e.g. as provided in a Variant Call Format (VCF) file. The multiple alignment is a matrix with genomes in rows and columns representing genomic offsets. The elements are either bases or gaps. We call a column consisting of identical bases and lacking any gaps a *uniform* column. Any other column is a *polymorphic* column. Figure 1 illustrates a multiply-aligned collection of haplotypes and the concatenated text *T*.

**Figure 1.**
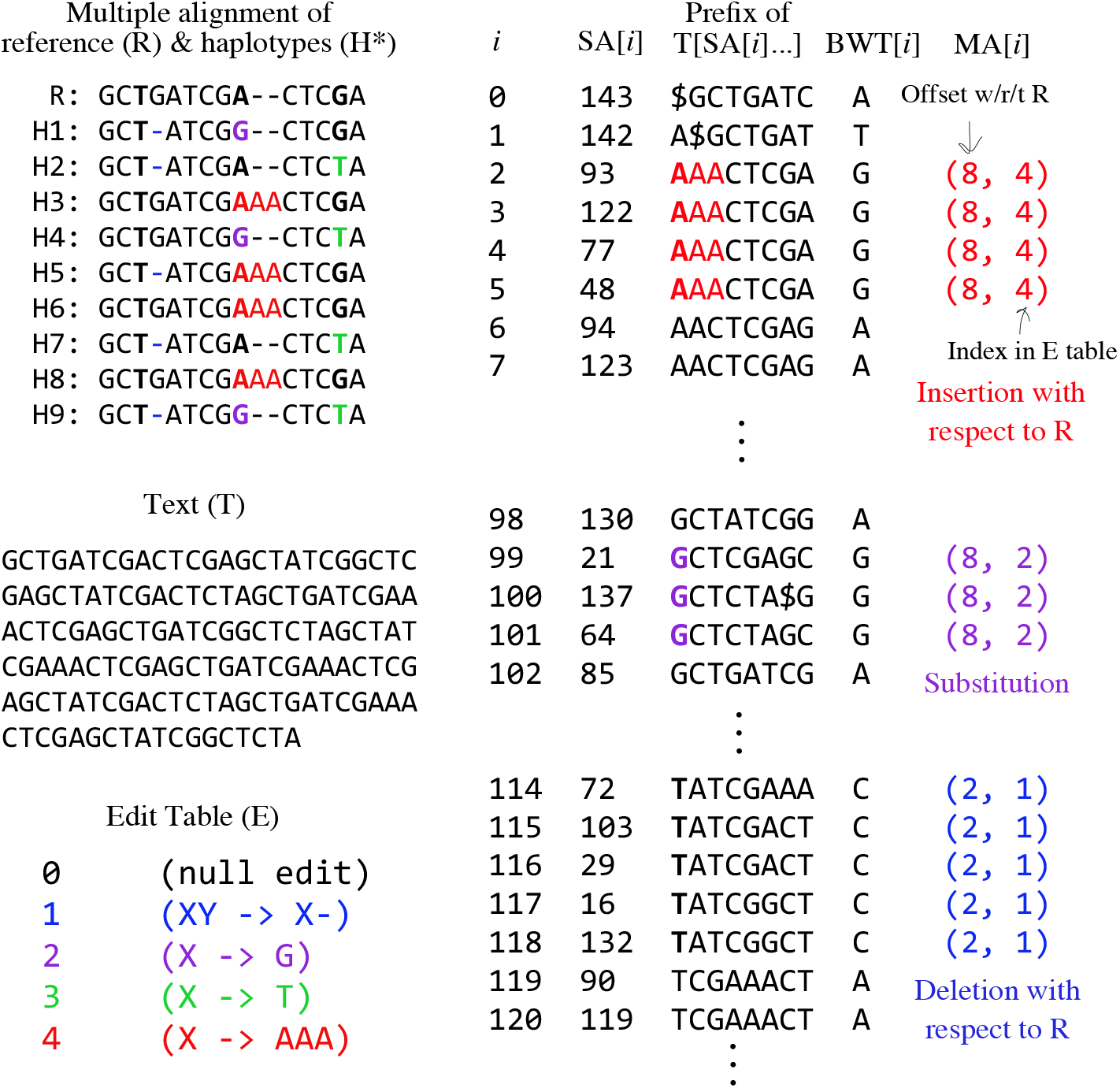
Top left: A multiple alignment for a collection of alternate haplotypes (`H1–H9`), and a reference sequence (`R`). Marked bases are in bold and alternate alleles are colored. Middle left: The text *T*, formed by concatenating rows of the multiple alignment (eliding gaps). Bottom left: The edit table *E*, with alternate-allele coloring. Right: A partial illustration of the marker array in relation to `SA`, the relevant suffixes themselves (truncated to fit), and the `BWT`. Colors and bolding highlight where marked bases and alternate alleles end up in the suffixes.

### Marker Array

Let the “marker array” `M` be an array parallel to the concatenated sequence *T* marking positions that fall in a polymorphic column in the multiple alignment. Each element of `M` is a tuple recording the offset *i* with respect to *T*_0_ where the polymorphism begins, as well as the edit operation describing how the sequence differs from the reference at this locus. Distinct edit operations are given distinct integer identifiers, which are decoded using a separate table *E*. Identifier 0 is the null operation, denoting that the reference allele appears without edits. For example, say *E* = {1 : *X* → *C*}, where *X* → *C* denotes a substitution that replaces the reference base with C. Then a marker array record *m* = (500, 0) marks a locus with no edit with respect to reference position *T*_0_ [500]. A record *m*′ = (500,1) denotes that a substitution changes that base at *T*_0_[500] to a *C*. An example is shown in Figure 1 (bottom left).

Consecutive substitutions are collapsed into a single edit in the *E* table. Insertions and deletions (“indels” for short) are treated somewhat differently; the offset carrying the “mark” is the one just preceding the indel (just to its left) in the multiple alignment. Importantly, the mark covers exactly one position in the genome, even if the insertion/deletion spans many bases. The marked position must come to the left of the indel to ensure that suffixes starting at the marked position include the allele itself. In the multiple alignment in Figure 1 (top left), for example, a deletion with respect to R occurs in the fourth-from-left column, but the marked position is in the third-from-left column.

The marker array `MA` is a permutation of `M` such that marks appear in suffix-rank order:

#### ▶ Definition 1.

*The* marker array `MA` *is the mapping such that* `MA`[*i*] = `M`[`SA`[*i*]].

Thanks to suffix-rank order, identical `M`[*i*]’s are often grouped into runs in `MA`, as seen in Figure 1 (right).

A *marker query* for pattern string *q* returns all *m* ∈ `M` overlapped by an occurrence of *q* in *T*. We can begin to answer this query using backward search (Equation 1) with *P* = *q*, giving the maximal `SA` range [*i*..*j*] such that *q* is a prefix of the suffixes in the range. Having computed [*i*..*j*], we know that {`MA`[*i*], …, `MA`[*j*]} contain markers overlapped by *q*’s leftmost character. To recover the markers overlapped by the rest of *q*’s characters, two approaches can be considered, detailed in the following subsections. The *FL* approach recovers the overlapped markers in a straightforward way but uses *O*(|*q*| · `occ`) time, where occ is the number of times *q* occurs in *T*. The heuristic backward-search approach requires only *O*(|*q*|) time but is not fully sensitive, i.e. it can miss some overlaps.

### *FL* **approach**

Say [*i*..*j*] is the maximal `SA` range such that all rows have *q* as a prefix. We can perform a sequence of FL steps, starting from each row *x* ∈ [*i*..*j*]. An FL step is the inverse of an LF step. That is, if we write an LF mapping step in terms of a rank query

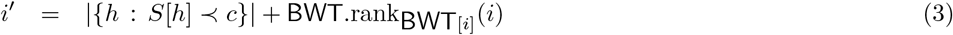

where *S*.rank_*c*_(*i*) denotes the number of occurrences of *c* in *S* up to but not including offset *i*, then an FL step inverts this using a select query

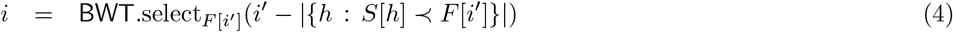

where *S*.select_*c*_(*i*) returns the offset of the *i* + 1^th^ occurrence of *c* in *S*, i.e. the *c* of rank *i*. Whereas LF takes a leftward step with respect to *T*, FL takes a rightward step.

By starting in each row *x* ∈ [*i*..*j*] and performing a sequence of |*q*| – 1 FL steps for each, we can visit each offset of *T* overlapped by an occurrence of *q*. Checking `MA`[*k*] at each step, where *k* is the current row, tells us which marker is overlapped, if any. This is slow in practice, both because it requires *O*(|*q*|(*j* – *i* + 1)) FL steps in total, and because each step requires a select query, which is more costly in practice than a rank query.

### Heuristic backward-search approach with smearing

Say we perform a backward search starting with the rightmost character of *q*. At each step we are considering a range [*i*..*j*] of `SA` having a suffix of *q* as a prefix. Using *i* and *j*, we can query `MA`[*i*..*j*]. However, this tells us instances where a *suffix* of *q* overlaps a marker, whereas our goal is to find where the *whole* query *q* overlaps a marker. If we report overlaps involving trivially short suffixes of *q*, many would be false positives. We propose to allow but reduce the number of such false positives by augmenting `MA`:

#### ▶ Definition 2.

*The augmented marker array* `MA`^*w*^ *is a multimap such that* `MA`^*w*^[*i*] = [`M`[`SA`[*i*]], `M`[`SA`[*i*] + 1],…, `M`[`SA`[*i*] + *w*]]

That is, `MA`^*w*^[*i*] is a (possibly empty) list containing markers overlapping any of the positions *T*[*i*..*i* + *w*]. We call this a “smeared” marker array, since the marks are extended (smeared) to the left by *w* additional positions. Note that a length-*w* extension can overlap one or more other marked variants to the left. For this reason, `MA`^*w*^ must be a multimap, i.e. it might associate up to *w* markers with a given position.

Using `MA`^*w*^, we adjust the backward-search strategy so that instead of querying `MA` at each step, we query `MA`^*w*^ every *w* steps. If *w* is large enough — e.g. longer than the length at which we see random-chance matches — we can avoid many false positives. More space is required to represent `MA`^*w*^ compared to `MA` since it is less sparse. However, we expect `MA`^*w*^ to remain run-length compressible for the same reason that `MA` is.

### Genotyping a read

Given a sequencing read, we would like to extract as much genotype information as possible while minimizing computational cost and false-positive genotype evidence. Here we give a heuristic algorithm (Algorithm 1) that handles entire sequencing reads, querying `MA`^*w*^ during the backward-search process as proposed in the previous section. The algorithm proceeds right to left through the read, growing the match by one character if possible. When we can no longer grow the match (i.e. the range [*i*..*j*] becomes empty), we reset the range to the all-inclusive range [0..|*T*| – 1] and restart the matching process at the next character. We use the term “extension” to refer to a consecutive sequence of steps that successfully extend a match. Note that this is a heuristic algorithm that does not exhaustively find all half-MEMs between the read and the index, as the MONI algorithm does [25].

#### Algorithm 1

Simplified version of `rowbowt` algorithm for matching query string *q* and compiling genotype evidence. *C* is an array such that *C*[*c*] equals |{*h* : *T*[*h*] ≺ *c*}|, where *T* is the original text. *N_h_* is the number of haplotypes in the index. Pseudocode for TallyEvidence and FinalizeEvidence are not given, but the heuristics they use to further filter genotype evidence is described in the text.

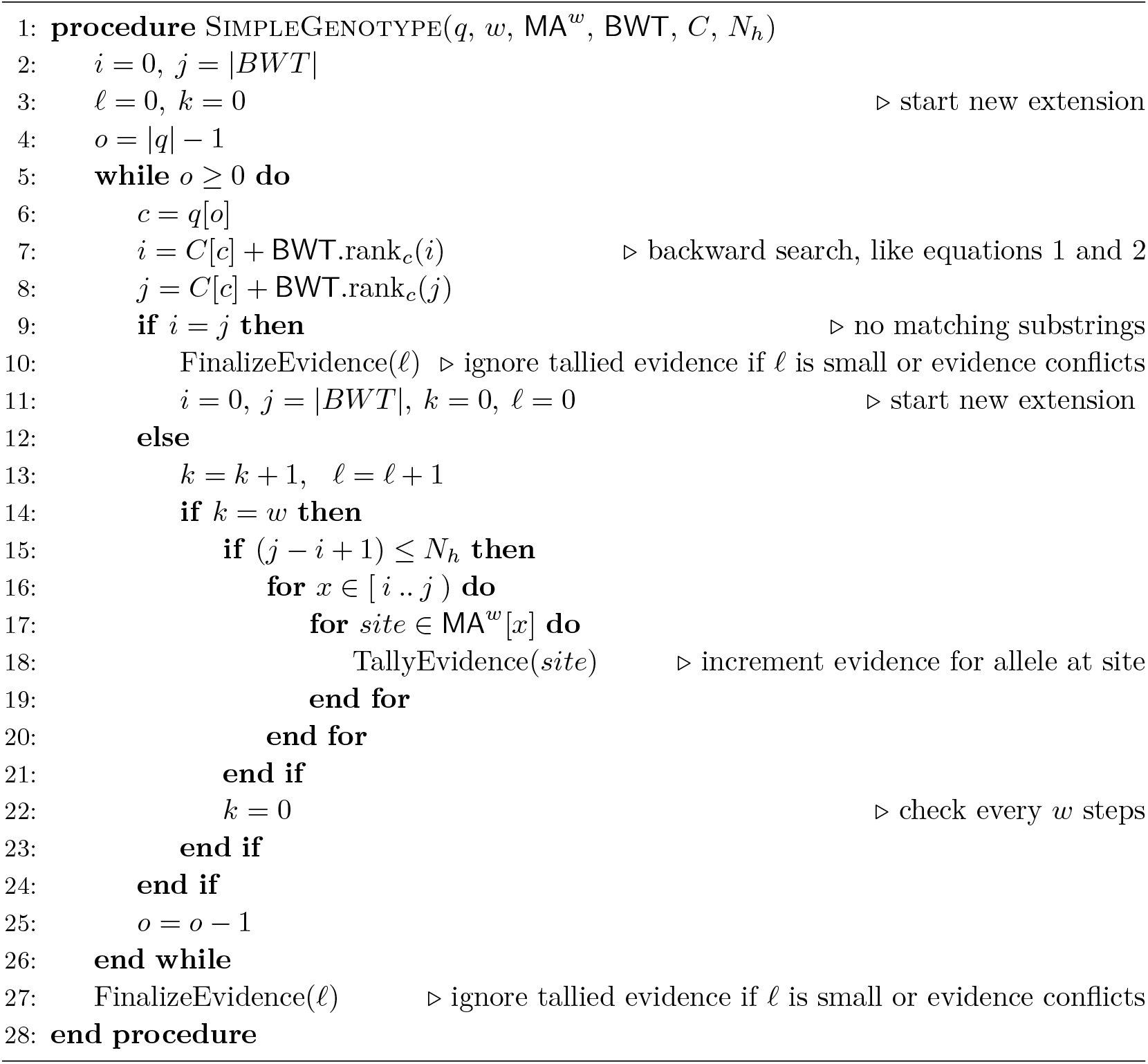

As discussed above, the algorithm only checks the marker array every *w* steps (line 14). As an additional filter, the algorithm only performs a marker-array query when the current suffix-array range size no larger than the number of haplotypes in the index (*N_h_*). A range exceeding that size indicates that we are seeing more than one distinct match in at least one haplotype, meaning that the evidence is ambiguous.

The algorithm tallies evidence as it goes (line 18), but might later choose to ignore that evidence if certain conditions are not satisfied (lines 10 and 27). For example, if the evidence has a conflict — i.e. one match indicates a reference allele at a site but another match found during the same extension finds an alternate allele at that site — then all the evidence is discarded for that extension. Similarly, evidence from an extension is discarded if the tallied sites span multiple chromosomes. Finally, evidence from extensions failing to match at least 80 bp of the read (adjustable with `--min-seed-length` option) is discarded.

We employ other heuristics to minimize mapping time not shown in Algorithm 1. For instance, we avoid wasted effort spent querying the wrong read strand. Specifically: `rowbowt` randomly selects an initial strand of the read to investigate: forward or reverse complement. If an extension from this strand meets the minimum seed-length threshold (80 by default), then the other strand is not considered and analysis of the read is complete. Otherwise, rowbowt then goes on to examine the opposite strand of the read.

### Sparse marker encoding

We encode the sparse arrays `M, MA` and `MA`^*w*^ in the following way. Say that array *A* consists of empty and non-empty elements. We consider *A*’s non-empty elements as falling into one of *x* maximal runs of identical (and non-empty) elements. Our sparse encoding for *A* consists of three structures. *S* is a length-|*A*| bit vector with 1s at the positions where a run of identical entries in *A* begins, and 0s elsewhere. *E* is a similar bit vector marking the last position of each of the *x* runs. (This variable *E* is distinct from the *E* table defined above in “Marker Array.”) To find whether an element *A*[*i*] is non-empty, we can ask whether we are between two such marks; that is, *A*[*i*] is non-empty if and only if *S*.rank_1_(*i* + 1) > *E*.rank_1_(*i*).

*X* is a length-*x* array containing the element that is repeated in each of *A*’s non-empty runs, in the order they appear in *A*. If *A*[*i*] is not empty, the element appearing there is given by *X*[*E*.rank_1_(*i*)].

When encoding `M` or `MA`, the elements of *X* are simply tuples. A complication exists for `MA`^*w*^, since elements are lists of up to *w* tuples. In this case, we keep an additional bit-vector *B* of size |*X*| where 1s denote left-hand boundaries in *X* that correspond to runs in *A*. *E* and *B* can be used together to access an element in *A* (Algorithm 2).

#### Algorithm 2

Access the contents of *A*[*i*] in the case where entries of *A* can be lists

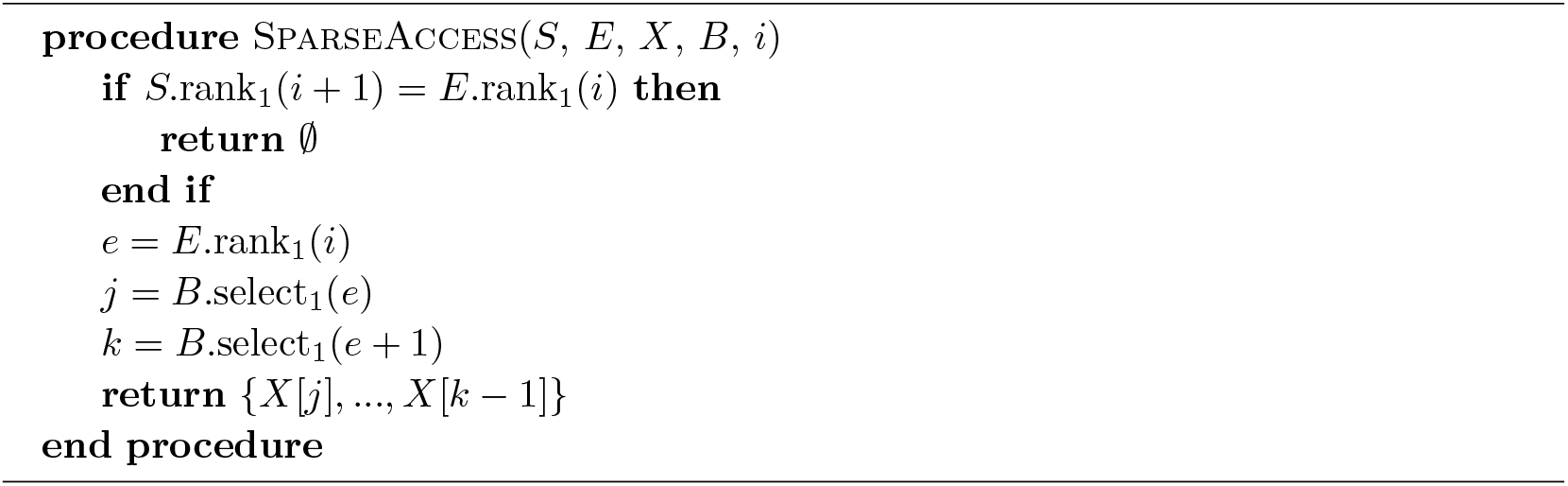

### Extracting markers from VCF

A Variant Calling Format (VCF) [9] file is used to encode a collection of haplotypes with the variants arranged in order according to a reference genome. In the case of human and other diploid genomes, haplotypes are grouped as pairs. We refer to such a collection of haplotypes as a “panel” and a single haplotype as a “panelist.” An VCF entry encodes a variant as a tuple consisting of a chromosome, offset, the allele found in the reference, the alternate allele found in one or more panelists, and a sequence of flags indicating whether each panelist has the reference or alternate version. We start from a VCF file to determine how to populate the marker arrays `M`, `MA` and/or `MA`^*w*^, as well as the edit array *E*.

A single element of `M` is a tuple (*r, e*), where *r* is an offset in *T*_0_ and *e* is the edit operation describing how the sequence differs from the reference. As a practical matter, we represent these tuples in a different way that more closely resembles the corresponding VCF records. Specifically, a marker is encoded in a 64-bit word divided into three fields. First, a 12 bit field identifies the chromosome containing the marker. The chromosome ordering is given at the beginning of the VCF file in the “header” section. For example, if “chr1” is the first chromosome in the header, then this chromosome is encoded as `0×000` (using hexadecimal), and if “chr2” is the second chromosome, it is encoded as `0×001`. Second is a 54 bit field encoding the marker’s offset within the chromosome. Third is a 4-bit field storing which version of the variant is present, with 0 indicating the reference allele, 1 indicating the 1st alternate allele, 2 the second alternate allele, etc. This 64-bit representation allows for compact storage of markers and easier random access to the marker array.

### Diploid genotyping

In a diploid genome, it is possible for both alleles to occur, i.e. for the genotype to be heterozygous. We use an existing approach [21] to compute genotype likelihoods considering all possible diploid genotypes: homozygous reference (2 reference alleles), homozygous alternate (2 alternate alleles), or heterozygous (1 reference, 1 alternate). Let *g* ∈ {0, 1, 2} denote the number of reference-allele copies at the marked site; e.g. *g* = 1 corresponds to a heterozygous site and *g* = 2 to a homozygous reference site. Let *l* be the number of times the reference allele was observed in the reads overlapping a particular marked site and let *k* be the count of all alleles (reference or alternate) observed. Let *ϵ* be the sequencing error rate. We calculate the genotype likelihood as follows, adopting equation 2 of [21] while setting the ploidy to 2 and adopting a global rather than a per-base error rate:

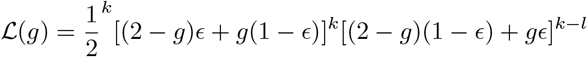

To choose the most likely genotype *g_max_*, we compute:

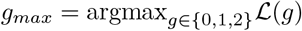

By default, `rowbowt` uses *ϵ* = 0.01.

### Implementation details

The code for constructing the marker array is implemented in the `pfbwt-f` spoftware package, with repository at https://github.com/alshai/pfbwt-f. This repository also contains an efficient implementation of the prefix-free-parse BWT construction algorithm [19]. This software is written in C++17, uses the open-source MIT license, and builds on the Succinct Data Structure Library (SDSL) v3.0 [16].

For querying the marker array, we use the `rowbowt` implementation at https://github.com/alshai/rowbowt. This repository contains the open source C++17 implementation of `rowbowt`, distributed under the MIT license. It is also a library, containing algorithms for building and querying indexes containing various structures discussed here, including the run-sampled suffix array, marker array, and others.

## 4 Results

We evaluated the efficiency and accuracy of our marker-array method for compiling genotype evidence. We first generated multiple series of `rowbowt` indexes covering various settings for three parameters: the window size *w* for the smeared marker array `MA`^*w*^, the number of haplotypes indexed, and the minimum allele frequency for marked alleles. The `rowbowt` index consisted of three components: the run-length encoded BWT, the run-sampled suffix array, and the marker array. While we built the sampled suffix array for these experiments, the standard marker-array-based method in `rowbowt` does not require this array.

We generated indexes for collections of 200, 400, 800, or 1000 human chromosome-21 haplotypes from the 1000 Genomes Phase 3 reference panel [2] based on the *GRCh37* reference. We generated two sets of indexes: one where the marker array marks all polymorphic sites regardless of frequency (denoted “*AF* > 0”), and another where the marker array marks only those sites where the less common allele occurs in greater than 1% of haplotypes, i.e. has allele frequency over 1% (denoted “*AF* > 0.01”). In all cases, the marker array window size *w* was set to 19. Each haplotype collection was drawn from a random subset of 500 individuals from the 1000 Genomes Phase 3 panel. The *AF* > 0 panel of 500 haplotypes contained 1, 097, 388 polymorphic sites. The *AF* > 0.01 panel of the same haplotypes contained 193, 438 polymorphic sites with allele frequency over 1%. We also included the GRCh37 reference sequence, consisting of all reference alleles, in each collection, corresponding to the reference sequence called *T*_0_ above.

We generated a series of indexes with window size *w* ∈ {13, 15, 17, 19, 21, 23, 25}. We generated two such series: one with no minimum allele frequency (*AF* > 0) and another with a 1% minimum frequency (*AF* > 0.01). Each index was over the same set of 100 haplotypes.

### Index size

We measured the size of the three main components of the rowbowt index: the augmented marker array, the run-length encoded BWT (RLE BWT) [3] and the run-sampled suffix array (“*r*-index SA”) [14]. Figure 2 plots this measurement for collections of 200, 400, 800 and 1,000 haplotypes for both *AF* > 0 and *AF* > 0.01. All grow linearly with the number haplotypes grows, as expected. For *AF* > 0, the augmented marker array is consistently larger than the run-sampled suffix array (“*r*-index SA”). For *AF* > 0.01, the augmented marker array is much smaller, approaching the size of the RLE BWT. The *AF* > 0 array is larger because it contains polymorphic sites with infrequent alleles; about 85% of the marked sites in the *AF* > 0 array have allele frequency under 1%. Further, rare alleles are less likely to form long runs in the augmented marker array, negatively affecting run-length compression.

**Figure 2.**
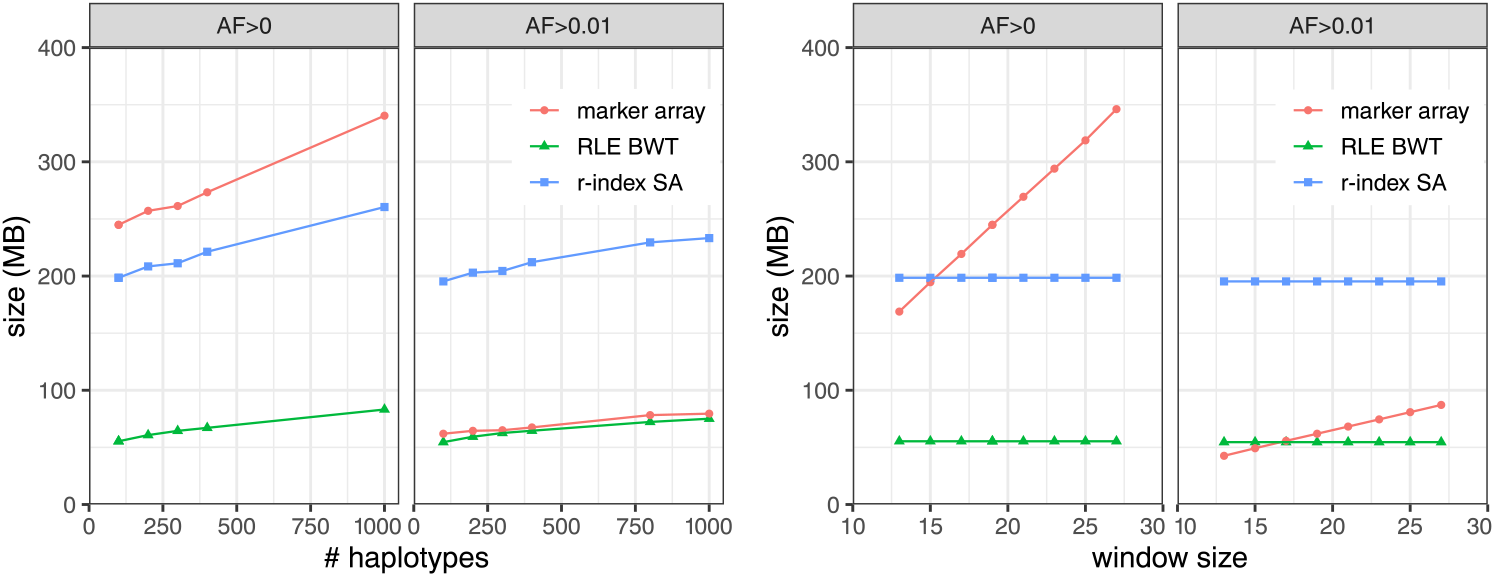
Left: Size of `rowbowt` data structures as a function of the number of haplotypes indexed, and with *w* = 19. “Marker array” refers to the augmented marker array, `MA`*^w^*. Right: Size of rowbowt data structures as a function of the “smearing” window size *w*, with number of haplotypes fixed at 100. Separate results are shown for when there is no minimum allowed allele frequency (*AF* > 0) and when the minimum frequency is 1% (*AF* > 0.01).

In the right portion of Figure 2, the RLE BWT and *r*-index SA have constant size because the *w* parameter does not affect those data structures. In the left portion of Figure 2, showing size as a function of number of haplotypes, the augmented marker array is almost always larger than the *r*-index SA for *AF* > 0 as opposed to *AF* > 0.01, except at *w* = 13. The slope of the array size is smaller for *AF* > 0.01 than for *AF* > 0.

Overall, both the value of *w* and the number of haplotypes in the index cause the augmented marker array to increase in size, but the inclusion of rare alleles (< 1% allele frequency) has the largest effect on its size.

### Query time

We next measured query time for the augmented marker array strategy versus the locate-query strategy which uses the run-sampled suffix array. 150bp simulated reads of 25× coverage were generated for one haplotype of HG01498, an individual that is part of the 1000-Genomes panel, but which we excluded from all our indexes. We simulated reads using Mason 2 `mason_simulator` [24] with default options.

In the case of the marker-array strategy, we measured the time required to analyze the reads using the algorithm described above in “Genotyping a read.” In the case of the locate-query strategy, the `MA`^*w*^ query was replaced with a two-step process that first performed a locate query with respect to the run-sampled suffix array, then performed a lookup in the `M` array. To enable this mode, we further augmented the *r*-index with a representation of `M` using the sparse encoding described above. To emphasize: the `rowbowt` strategy does not require the run-sampled suffix array or the `M` array; the `MA`^*w*^ effectively replaces them both.

We repeatedly sampled 10,000 simulated reads and recorded the mean query time over 10 trials. As seen in Figure 3 the augmented marker-array method (labeled “marker”) was consistently faster than locate method. This was true for all allele frequencies and window sizes tested. We found that the effect of *w* and allele-frequency cutoff was more pronounced with the larger reference panel *AF* > 0. For the smaller panel (*AF* > 0.01), query time was mostly invariant to both window size and allele frequency.

**Figure 3.**
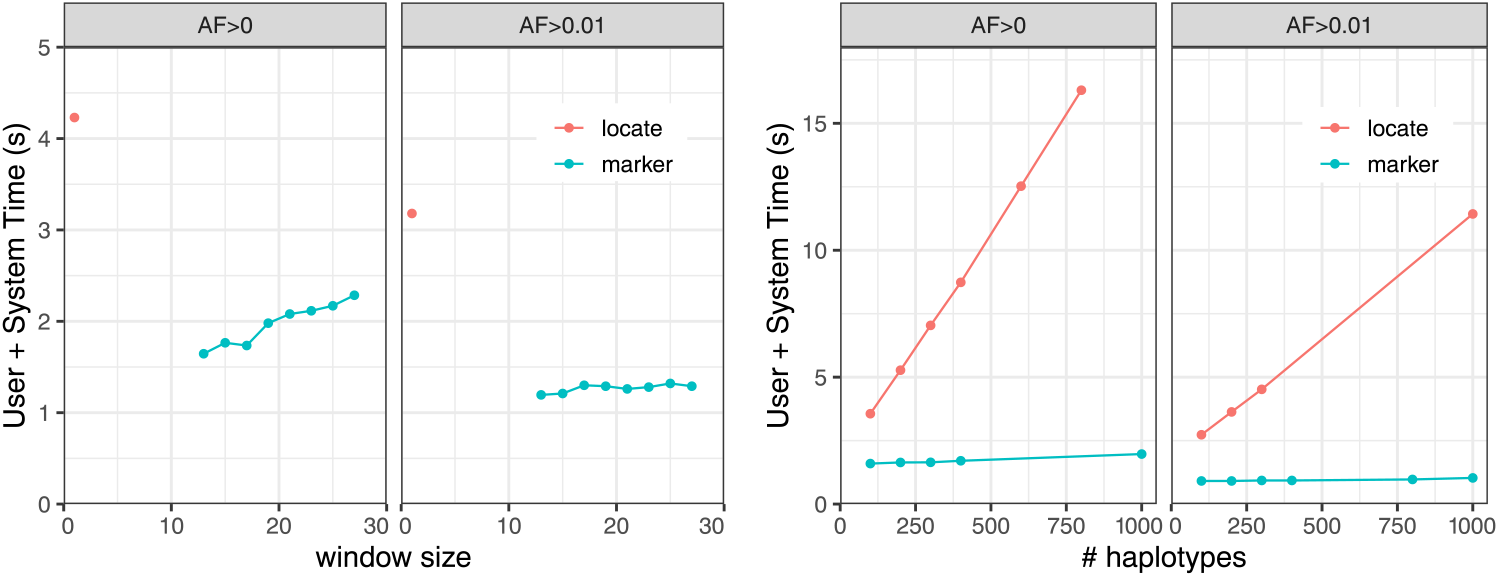
Mean time over 10 trials of aligning 10,000 simulated reads from HG01498 against the augmented marker array (marker) and the *r*-index suffix array (locate). Experiments are repeated for marker collections including all alleles (*AF* > 0) and for alleles having frequency at least 1% (*AF* > 0.01). Left: The experiment is repeated for various window sizes *w*, and for 100 haplotypes. Right: The experiment is repeated for different numbers of indexed haplotypes, with *w* = 19.

### Genotyping accuracy

We next measured the accuracy of the genotype information gathered using the augmented marker array. We simulated sequencing reads from one haplotype of HG01498 to an average depth of 25-fold coverage. Individual HG01498 was excluded from the indexes. As our “truth” set for evaluation, we use the variant calls in the 1000 Genomes project callset for the same haplotype we simulated from. For simplicity, this experiment treats the genome as haploid. Experiments in the next section will account for the diploid nature of human genomes.

A single marked site can have conflicting evidence, due, for instance, to mismapped reads or sequencing errors. For this evaluation, we make calls simply by finding the frequently observed allele at the site. We ignore any instances of alleles other than the ones noted in the VCF file as the reference and alternate alleles. If the reference and alternate alleles have equal evidence, the reference allele is called.

We calculate precision and recall according to the following formulas. Here, the positive class consists of marked sites that truly have the alternate allele, while the negative class consists of marked sites that truly have the reference allele. We measure:

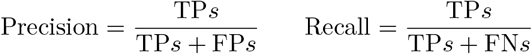

Where TP stands for True Positive, FN stands for False Negative, etc.

Figure 4 shows precision and recall with respect to the number of haplotypes in the index and the minimum allele frequency of the haplotype collection. We observed that the *AF* > 0.01 indexes generally had better precision compared to the *AF* > 0 indexes, though at the expense of recall. Precision and recall generally improve with the addition of more haplotypes to the index. The augmented marker array has similar recall to the locate procedure across all haplotype sizes at the loss of precision. When rare variants are removed from the index (*AF >* 0.01), the gap in precision between the marker array and the locate procedure lessens. This mild (less than 0.1%) loss of precision is expected since algorithm described above in “Genotyping reads” is still prone to some false positives in the earlier part of the extension process.

**Figure 4.**
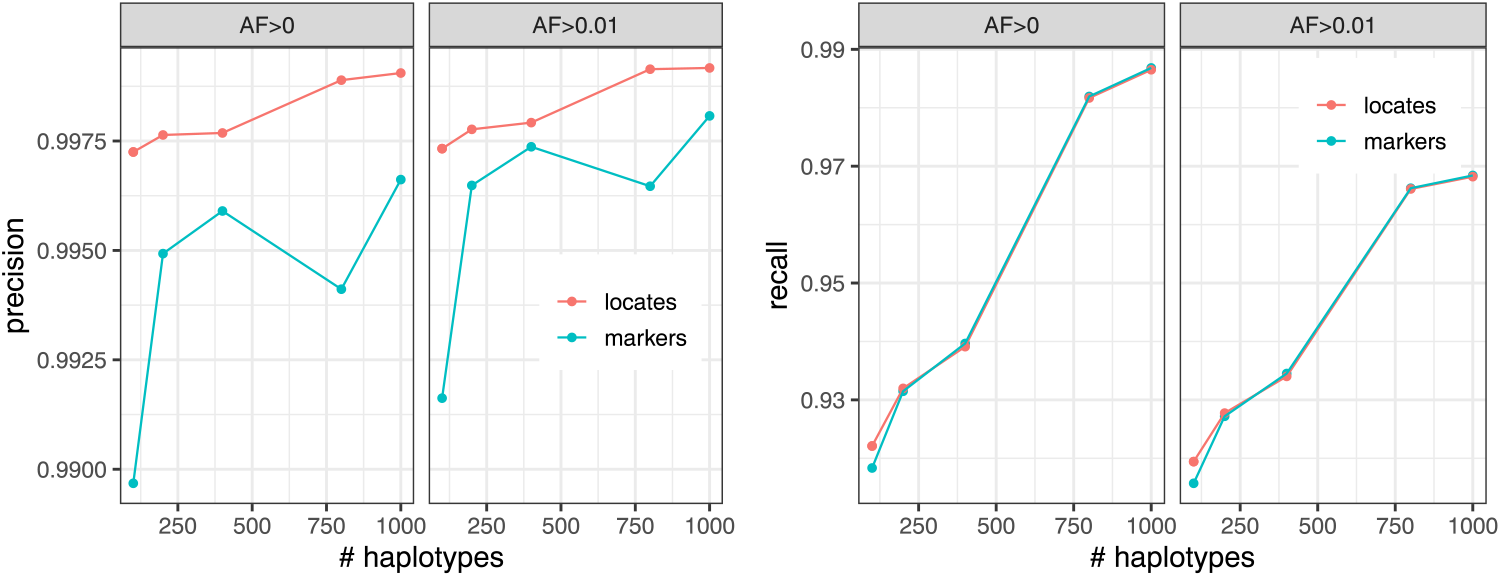
Precision (left) and recall (right) of the calls made when querying 25× simulated reads from HG01498 against the augmented marker array (marker) and the *r*-index (locate). Stratified by minimum allowed allele frequency (AF) in the index.

### Diploid genotyping assessment

To assess diploid genotyping accuracy, we used data from the Human Genome Structural Variation Consortium (HGSVC) [6, 12]. The HGSVC called both simple and complex genetic variants across a panel of 64 human genomes. Calls were made with respect to the GRCh38 primary assembly [26]. For input reads, we subsampled reads from a 30-fold average coverage PCR-free read set provided by the 1000 Genomes Project [2] (accession SRR622457). To create more challenging scenarios for the genotypers, we randomly subsampled the read sets to make smaller datasets of 0.01, 0.05, 0.1, 0.5, 1, 2 and 5-fold average coverage.

We assessed three genotyping methods. The first (`bowtie2_bcftools`) used Bowtie 2 [20] v2.4.2 to align reads to a standard linear reference genome, then used BCFtools v1.13 to call variants (i.e. genotypes) at the marked sites [21]. The second method used the graph-based genotyper `bayestyper` v1.5. The third was `rowbowt`.

Prior to applying `bayestyper`, we built a `bayestyper`-compatible VCF file containing all relevant variants from the HGSVC haplotype panel. For `rowbowt`, we created a `rowbowt` index from the genomes in the HGSVC haplotype panel. In both cases we excluded NA12878’s haplotypes from the panel prior to building the index.

When evaluating, we stratified variants by complexity: the “snp” category includes single-nucleotide substitutions, “indel” includes indels no more than 50bp long, and “sv” includes insertion or deletions longer than 50bp, and “all” includes all variant types. More complex structural variants like inversions and chromosomal rearrangements are ignored.

We analyzed the accuracy of `rowbowt`’s diploid genotypes in two ways. First we considered allele-by-allele precision and recall, considering the alternate (ALT) allele calls to be the positive class. Specifically, every diploid genotype called by a method is considered as a pair of individual allele calls. If a given allele call is an alternate (ALT) allele and there is at least one ALT allele present in the true diploid genotype at that site, it counts as a true positive (TP). If the given allele is a reference allele (REF) and there is at least one REF allele in the true diploid genotype, this is a true negative (TN). If the given allele is an ALT but the true genotype is homozygous REF, we count it as a false positive (FP). Finally, if the given allele is REF but the true genotype is homozygous ALT, this is a a false negative (FN).

Second, we considered precision and recall with respect to sites that were either truly heterozygous or called heterozygous. If a heterozygous call made by a method is truly heterozygous, this was counted as a true positive (TP). False positives, false negatives, and true negatives are defined accordingly.

As seen in Figure 5, `rowbowt`’s ALT and HET precision were generally the highest of all the methods across all variant categories, though `bayestyper` sometimes achieved higher ALT/HET precision for indels in the higher-coverage datasets. `rowbowt`’s ALT recall is also higher than the other methods, except for some of the lower coverage measurements in the “sv” category, where `bayestyper` achieved higher ALT recall.

**Figure 5.**
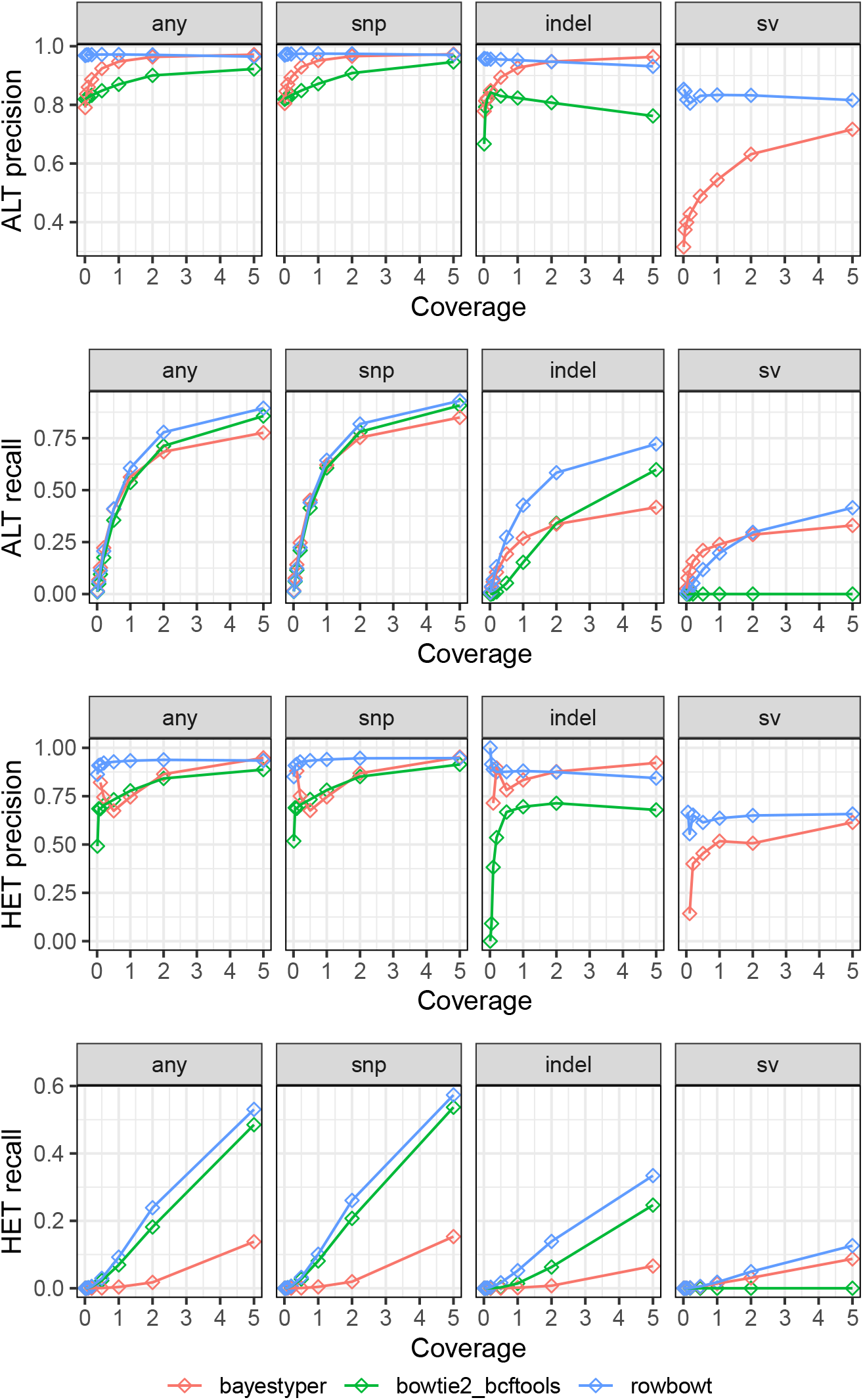
Precision and recall for the three tested genotyping methods, both at the level of individual alleles (ALT precision/recall) and at the level of heterozygous variants (HET precision/recall). Note that the bowtie2_bcftools approach is generally unable to align reads across variants in the “sv” category, leading to low precision and recall.

### Computational performance

We compared the time and memory usage of each genotyping method, dividing each method into Alignment and Genotyping phases. For each phase, we measured both the wall-clock time elapsed and maximum memory (“maximum resident set”) used. Both were measured with the Snakemake tool’s `benchmark` directive [22].

Since the tools operate somewhat differently, we define the “Alignment” step differently in each case. For `bowtie2_bcftools`, we define the Alignment step as the process of using `bowtie2` to align sequencing reads to the linear reference genome. For `rowbowt`, we defined the Alignment step as the process of using the `rb_markers` command to genotype the reads using the algorithm described in Methods. For `bayestyper`, we defined the Alignment step as the typical 3-step process of: (a) using the KMC3 software tool to count *k*-mers in the input reads, (b) using the bayestyper makeBloom command to convert *k*-mer counts to Bloom filters for each sample, and (c) using the `bayestyper cluster` command to identify variant clusters. 16 threads were used during the Alignment phase for `bowtie2_bcftools` and `rowbowt`, while 32 threads were used for bayestyper.

For `bowtie2_bcftools`, we define the Genotyping step as the process of using `bcftools` call to call variants from the BAM file output by `bowtie2`. For `rowbowt` We define the Genotyping phase as the process of running the `vc_from_markers.py` script on the output from `rb_markers`. For `bayestyper`, we define the genotyping step as the process of running the `bayesTyper genotype` command. The Genotype phases for both `bowtie2_bcftools` and `rowbowt` do not support multi-threading, so a single thread was used. For the `bayestyper` Genotype phase we used 16 threads.

Figure 6 shows the time taken and peak memory footprint for each method and each phase. We observed that `rowbowt` was consistently faster than the other methods, sometimes by a large margin. We also observed that while `rowbowt` has a higher memory footprint compared to the `bowtie2_bcftools` method, it uses substantially less memory than `bayestyper`, the other pangenome-based method.

**Figure 6.**
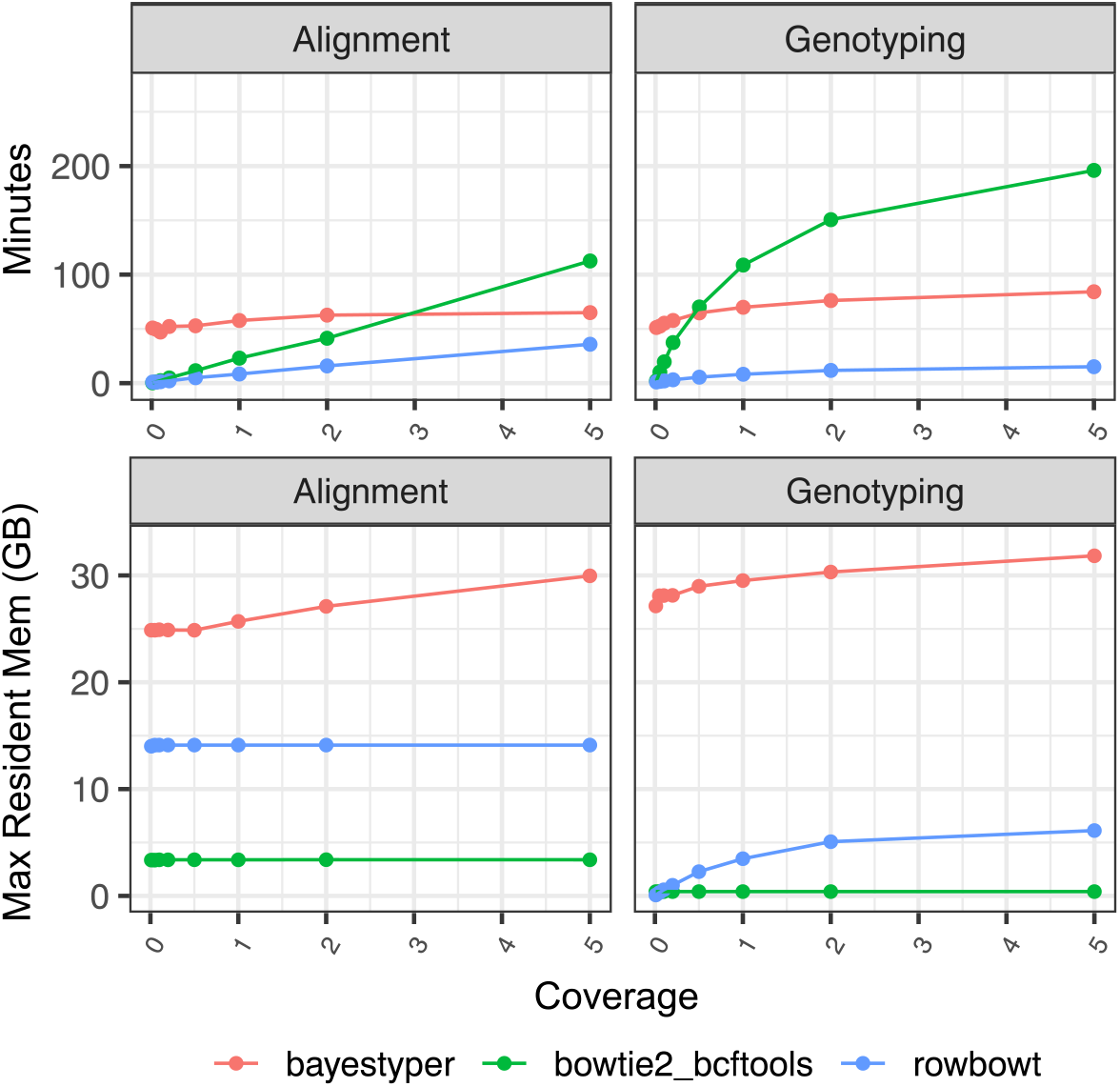
Time for each phase of the impute-first workflow for three methods of alignment/geno-typing

## 5 Discussion

We proposed a family of novel marker array structures, `M, MA` and `MA`^*w*^ that, together with a pangenome index like the *r*-index, allow for rapid and memory-efficient genotyping with respect to large pan-genome references. The augmented marker array is smaller and faster to query than the run-sampled suffix array — the usual way to establish where matches fall when querying a run-length compressed index — especially when we limit the set of markers to just alleles at frequency 1% or higher. We further showed that the augmented marker array can replace the sampled suffix array in simple genotyping experiments with moderate sacrifice of precision, and that a marker array based genotyping method outperforms the graph-based Bayestyper method.

Pan-genome indexes allow for rapid analysis of reads while avoiding reference bias. The indexes used in our experiments consisted of many (up to 65) haplotypes, with none having a higher priority over the others, except in the sense that results were expressed in terms of the standard reference. Our approach preserves all linkage disequilibrium information. This is in contrast to some graph-indexing approaches, which might consider all possible combinations of nearby alleles to be “valid,” even if most combinations never co-occur in nature.

While we examined only simple structural variants in the form of insertions and deletions longer than 50 bp, the genotyping method is readily extensible to more complex differences as well. Indeed, as long as we can mark the base or bases just to the left of the variant, we can mark any variant in a way that we can later genotype

The `rowbowt` method can lead to future methods that use information about genotypes to build a personalized reference genome, containing exactly the genotyped alleles. Alignment to a personalized reference have been shown previously to be the best way to avoid reference bias, even more effective than the best pangenome methods [7, 23].

## 6 Ackonwledgements

We thank Massimiliano Rossi and Travis Gagie for many helpful discussions. We thank Margaret Gagie for her careful editing.

Part of this research project was conducted using computational resources at the Maryland Advanced Research Computing Center (MARCC).

